# Alternative promoter usage modulates miRNA-guided translation inhibition of a m^6^A reader in phosphate starvation

**DOI:** 10.1101/2020.12.19.423605

**Authors:** Rodrigo Siqueira Reis, Jules Deforges, Joaquín Clúa, Yves Poirier

## Abstract

Alternative transcription start sites (TSSs) are widespread in eukaryotes. In plants, light, development and tissue regulate selective usage of several TSSs, producing transcripts with distinct 5′UTR as well as shorter protein isoforms with distinct subcellular localization or activity. However, the function of non-coding transcripts generated by alternative TSSs is still largely unknown. We show that phosphate availability regulates numerous alternative TSSs, including a non-coding alternative TSS (*ALT*_*ECT4*_) associated with *ECT4*, encoding a N^6^-methyladenosine (m^6^A) reader. We found that *ECT4* harbors a cleavage-resistant miR826b target site at its 3’UTR, also present in *ALT*_*ECT4*_. In the absence of *ALT*_*ECT4*_, miR826b guides translation inhibition of *ECT4*. Phosphate deficiency triggers specific and robust expression of *ALT*_*ECT4*_, counteracting miR826b inhibition of its targets, including *ECT4*. The role of *ALT*_*ECT4*_ as a miR826b antagonist shows that it acts in *cis* to regulate translation of the m^6^A reader ECT4, and this function might be shared among other non-coding transcripts generated by alternative TSS.

## Introduction

Alternative TSS usage is an important mechanism influencing mRNA and protein diversity^1-5^. In humans, an average of four different transcription start sites (TSSs) are expressed per gene, many exhibiting tight tissue or cell-type regulation(Forrest et al., 2014; de Klerk and ‘t Hoen, 2015). Indeed, alternative transcription initiation and termination, rather than alternative splicing, are the main drivers of transcript isoform diversity in humans(Reyes and Huber, 2018). Alternative TSS is also common in plants and can affect the presence of regulatory elements found in mRNAs 5′ UTRs, often fine-tunning translation. For instance, blue light induces downstream TSS usage in two key light-inducible transcription factors, HY5 and HYH, producing shorter transcripts with higher translation efficiency as they lack inhibitory uORFs(Kurihara et al., 2018). In many cases, alternative TSS usage produces shorter protein sequences that display differential elements within their N-terminus, including signal sequences. Such isoforms play important role in light-dependent changes in subcellular localization of proteins, such as GLYK(Ushijima et al., 2017). GLYK is functional as a photorespiration enzyme in chloroplast, however, shade induces expression of an isoform that lacks the chloroplast signal, thus becoming cytoplasmic, which seems to play a role in adaptation to light fluctuations.

Alternative TSS usage can also produce transcripts isoforms that are significantly shorter than the annotated reference isoforms(Forrest et al., 2014; Wang et al., 2019), with many too short to code for functional proteins. However, the function of non-coding transcript isoforms originating from alternative TSS usage are poorly characterized and their functions remain unknown. Tight transcriptional regulation such non-coding alternative transcript hints towards a potential regulatory function on the main full-length transcript.

Here, we identified several Pi-specific alternative transcripts, including shorter transcripts lacking uORF and signal peptide sequences. Several of those transcripts lacked, either partially or fully, functional domain(s) found in the reference, full-length transcripts. The N^6^-methyladenosine (m^6^A) reader *ECT4* locus showed strong Pi-induction of an alternative TSS (*ALT*_*ECT4*_), lacking any ECT4-related functional domain. We then found that *ALT*_*ECT4*_ has a *cis*-regulatory function whereby it counteracts a miRNA-guided translation inhibition of ECT4. We further showed a role for the m^6^A pathway, including m^6^A writer and reader, in root development upon Pi deprivation. Our work revealed a complex network involving alternative TSS usage, miRNA regulation and m^6^A pathway, in which *ALT*_*ECT4*_ functions to modulate, in *cis*, translation of the main ECT4-coding transcript.

## Results and discussion

### Phosphate deficiency regulates alternative transcription start site

To identify transcripts with alternative TSS induced specifically by phosphate starvation, we initially applied JunctionSeq to a RNA-Seq dataset(Deforges et al., 2019a) that included plants grown under Pi-deficient condition or treated with various hormones. (Fig. 1a). JunctionSeq is designed to analyse differential usage of exonic regions(Hartley and Mullikin, 2016), which includes alternative TSS and splicing isoforms. This approach identified 294 differential transcripts associated with Pi deficiency, of which 259 (88%) were specific to this treatment (Fig. 1b and Table S1). Next, we analysed the same Pi-deficient samples for full-length RNAs (capped and polyadenylated) using TeloPrime(Cartolano et al., 2016) cDNAs sequenced using PacBio long-read (Fig. 1a). In total, 739 unique genes were associated with at least one alternative TSS regulated by Pi deficiency, of which 54 were also found in the JunctionSeq Pi-specific group (Fig. 1c and Table S1). As previously observed, several alternative transcripts were shorter at the 5’UTR but retaining the full-length coding sequence (CDS), while others had shorter CDS skipping putative signal peptide sequence or functional domain(s) (illustrated in Fig. 1d and Table S1). For instance, Pi deficiency induces expression of a long *RTNLB1* transcript harbouring six uORFs at its 5’UTR, which are absent in the alternative shorter transcript (Fig. 1d). Interestingly, the uORF-containing transcript was only expressed in Pi-deficient plants. Since uORFs are often repressors of translation(Srivastava et al., 2018), we compared the impact of the *RTNLB1* long 5’UTR with its shorter, alternative 5’UTR sequence, on translation. Protoplast expression and analysis of these 5’UTRs fused to a dual luciferase system, showed that fusion to *ALT*_*RTNLB1*_ 5’UTR results in much higher relative luminescence, compared with fusion to *RTNLB1* 5’UTR (Fig. S1), suggesting that the long *RTNLB1* 5’UTR harbors elements that inhibit protein translation under Pi deficiency.

**Figure 1.**
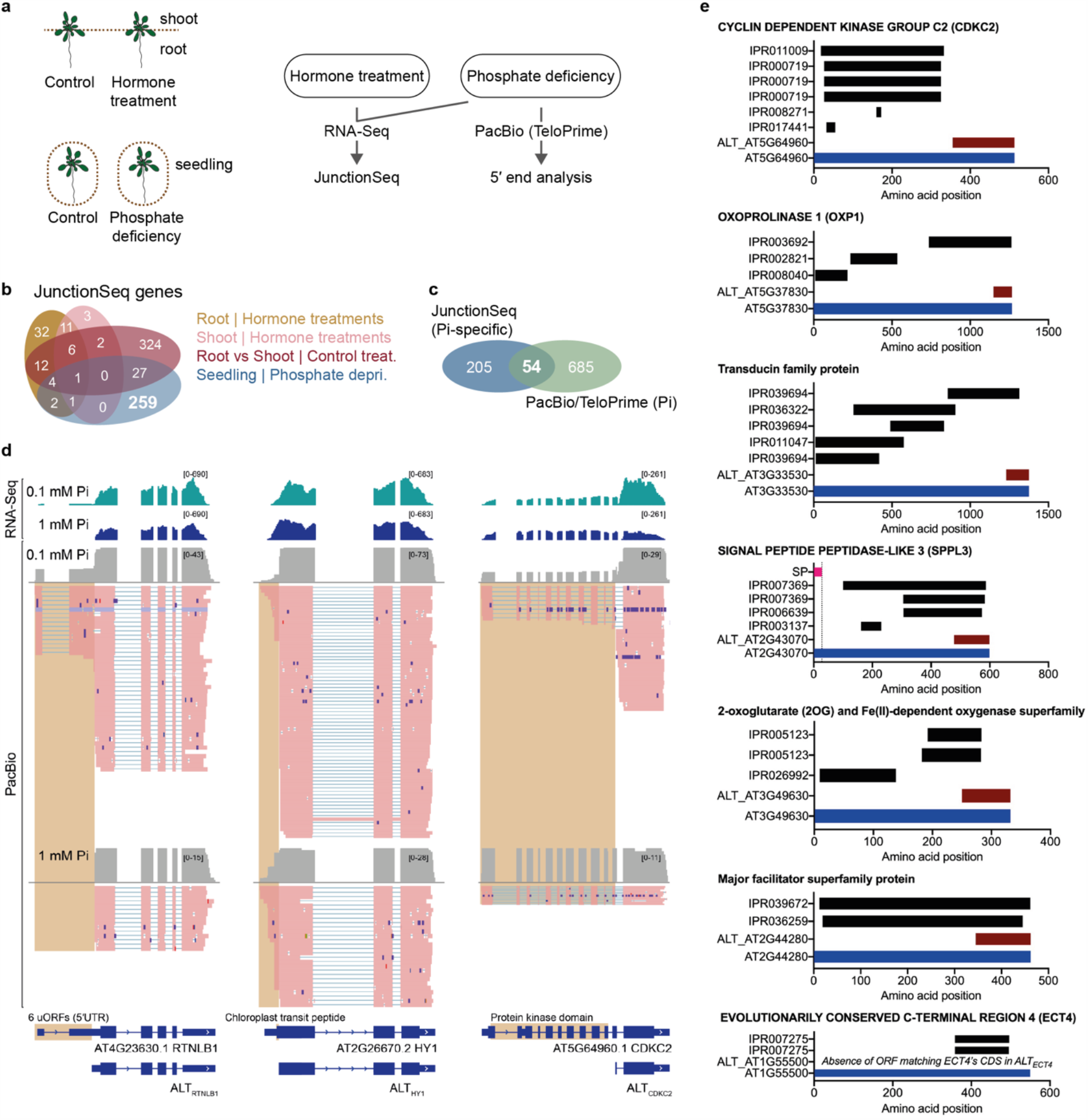
Phosphate deficiency regulates genome-wide alternative transcription start site. **a**, Experimental design of alternative TSS identification. Total RNA was extracted from root and shoot of hormone-treated 10-day old seedlings grown on solid MS medium, and Pi-deficient (1 mM vs 0.1 mM Pi) 7-day old seedlings grown in liquid MS medium. RNA-Seq data was produced for all samples and analyzed using JunctionSeq. PacBio long read data was produced from full-length cDNA samples (TeloPrime) of Pi-deficient plants. **b**, Venn diagram showing the number of genes significantly different (p-value<0.01) in JunctionSeq data. **c**, Venn diagram showing the number of genes significantly different in Pi-specific JunctionSeq (p-value<0.01) and PacBio (p-value<0.05) data. **d**, Selected genes exhibiting alternative TSS in Pi-deficient conditions. uORF, upstream open reading frame. Illumina RNA-Seq reads density graph (upper panel) and the PacBio reads for full length mRNAs (lower panel) are shown. **e**, Functional features associated with reference transcript-encoded amino acid sequence. Predicted domains using InterProScan depicted as IPR-number (black bar). SP, signal peptide (pink bar). Reference protein (locus name; blue bar) and alternative transcript-encoded amino acid sequence (ALT_locus name; red bar). Alternative transcript-encoded amino acid sequence was defined as the longest ORF ending at the reference transcript’s STOP codon, i.e. exhibiting the same (complete or partial) amino acid sequence as that found in the reference protein.

We found that adaptation to Pi deficiency might also involve changes in protein localization via signal peptide depletion, as a subset of alternative transcripts lack the sequence for signal peptide found in the reference transcript, while retaining the coding sequence for functional domain(s) (Fig 1d, e, Fig. S2 and Table S1). We also observed several alternative transcripts that lack (either fully or partially) sequence encoding for functional domain(s) found in the reference transcripts (Fig. 1e, Fig. S2 and Table S1). These transcripts might produce functionally different proteins and, in some cases, might function as non-coding RNAs. We then decided to focus our efforts on characterizing an alternative transcript derived from the *EVOLUTIONARY CONSERVED C-TERMINAL 4* (*ECT4; At1g55500*) locus that is likely to produce a non-coding RNA.

### Phosphate deficiency induces strong expression of a m^6^A reader-associated alternative promoter

*ECT4* encodes a highly conserved YTH-domain protein that functions as a N6-methyladenosine (m^6^A) reader(Arribas-Hernández et al., 2018). *ECT4* produce an alternative transcript, hereafter referred as *ALT*_*ECT4*_, initiated within *ECT4* 7^th^ intron, producing a 2-exon transcript of ∼550 nucleotides, including *ECT4* exon 8 and 9, thus lacking the YTH domain (Fig 1e, Fig. 2a and Table S2). We found 5 short ORFs within the *ALT*_*ECT4*_ transcript, but none in-frame with the full length *ECT4* CDS, suggesting that *ALT*_*ECT4*_ is either a noncoding RNA or produces ECT4-unrelated peptide(s) (Table S2). *ECT4* expression was slightly increased by Pi deficiency, whereas *ALT*_*ECT4*_ was strongly and specifically induced by this treatment (Fig. 2a and Fig. S2). This was further validated by RT-qPCR, evidencing a strong induction of *ALT*_*ECT4*_ in Pi-deficient roots (Fig. 2b). We then produced transgenic plants transformed with a construct containing the *ECT4* genomic sequence fused in-frame (*ECT4*’s STOP codon removed) with GFP, without *ECT4* upstream sequences (i.e. without 5’UTR and promoter), but including *ECT4* START codon. These plants showed enhanced free GFP expression under Pi-deficient condition, evidencing that expression of the *GFP* transcript is independent of the *ECT4* promoter in these lines, but relies on promoter elements within the coding region of *ECT4* (hereafter referred as *ALT*_*ECT4*_ promoter) (Fig. 2c). GUS expression driven by the *ECT4* promoter was similar in plants grown under Pi-sufficient and deficient conditions, being broad in shoots and mainly restricted to the vascular system in roots (Fig. 2d). In contrast, GUS expression from the *ALT*_*ECT4*_ promoter in plants grown under Pi-sufficient condition revealed a weak expression at the tip and base of leaf and cotyledon and no detectable expression in roots. Under Pi deficiency, GUS expression from the *ALT*_*ECT4*_ promoter expanded to the entire leaf, albeit stronger in vasculature, as well as in the vasculature of primary and lateral roots. In Pi-deficient plants, expression of GUS from the *ECT4* and *ALT*_*ECT4*_ promoters showed clear overlap in leaf and root vasculature.

**Figure 2.**
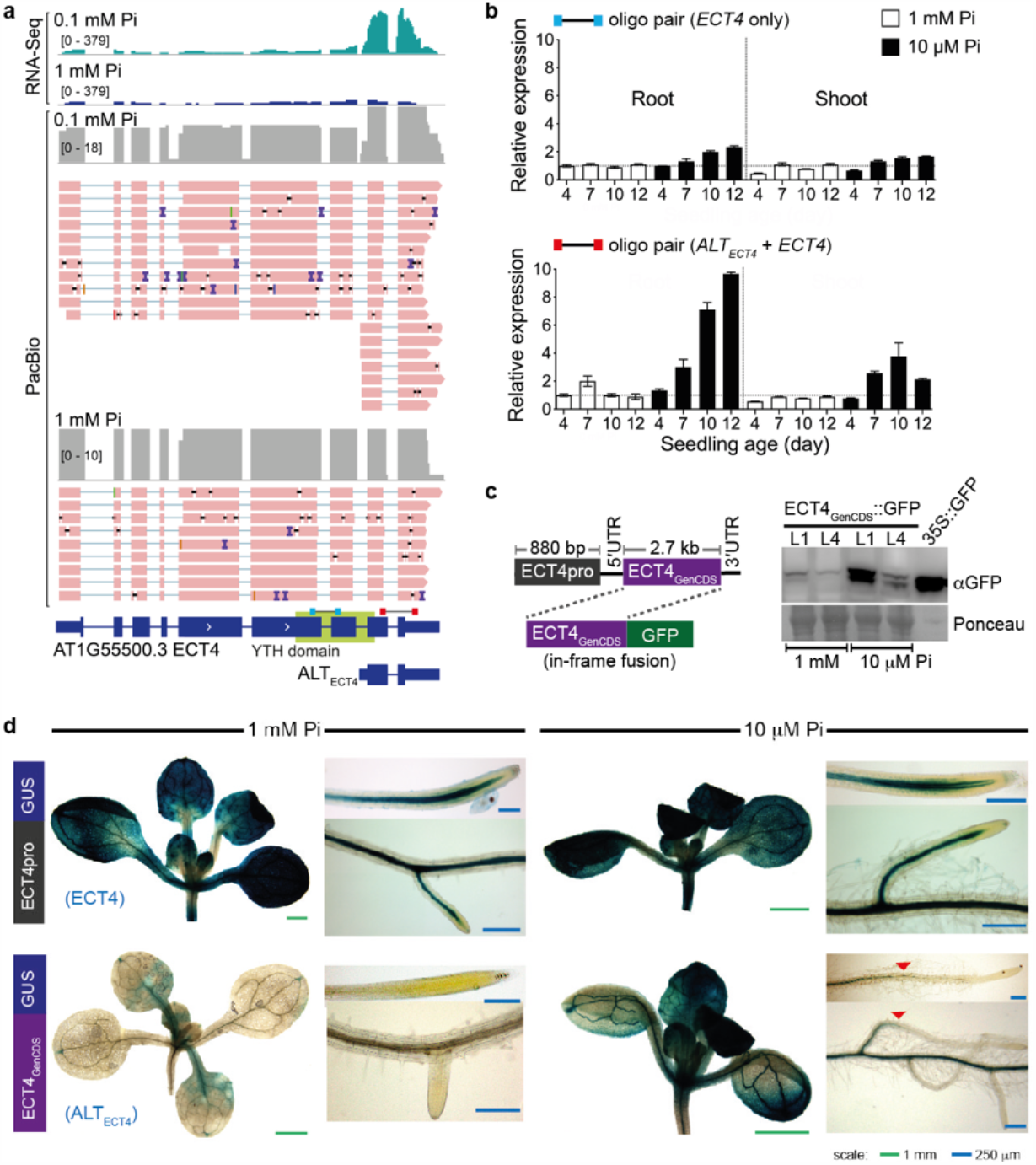
Pi deficiency induces alternative promoter selection associated with *ECT4*. **a**, Illumina RNA-Seq reads density graph (upper panel) and PacBio reads for full length mRNAs (lower panel) are shown for plants grown on Pi-sufficient and Pi-deficient media. *ALT*_*ECT4*_ sequence is depicted with *ECT4*’s annotation for coding sequence (thickest line; bottom panel). **b**, Time course of expression profile of *ECT4* and *ALT*_*ECT4*_ in plants grown on Pi-sufficient and Pi-deficient media. RT-qPCR using oligo pair (see **a**, bottom panel) specific to *ECT4* (top panel) or specific to both *ECT4* and *ALT*_*ECT4*_ (bottom panel). **c**, GFP expression driven by *ALT*_*ECT4*_ promoter. *ALT*_*ECT4*_ promoter includes ∼2.7 kb of the *ECT4* coding sequence (ECT4_GenCDS_), from the initiating ATG to the penultimate codon, fused in frame to GFP and expressed as transgenes. Free GFP control extracted from transgenic plants expressing *GFP* under the CaMV 35S promoter. **d**, Promoter-GUS staining profile using the 880 bp *ECT4* promoter region (*ECT4pro*) or the ALT_ECT4_ promoter (ECT4_GenCDS_) in tissues of plants grown on Pi-sufficient and Pi-deficient media. Red arrows in ALT_ECT4_ expression in 10 μM Pi mark the GUS staining borders. For all main panels: shoot (left), primary root (upper right) and lateral root(s) (lower right).

### The m^6^A pathway modulates primary root development

m^6^A is the major ribonucleotide modification in plants(Arribas-Hernández and Brodersen, 2020). In this pathway, proteins involved in m^6^A mRNA modification are referred to as writers (e.g. FIP37), effectors as readers (e.g. *ECT4*) and m^6^A removal is performed by erasers(Lim and Pawson, 2010) (partially depicted in Fig. 3a). To study a possible role for m^6^A modification in Pi homeostasis, we initially analysed null mutants for readers (i.e. *ECT2, ECT3* and *ECT4*), assuming genetic redundancy(Arribas-Hernández et al., 2018). *ect4* and *ect2ect3* mutants showed wild type primary root growth in Pi-deficient and sufficient conditions (Fig. 3b). While primary root growth of *ect2ect3ect4* triple mutant was only mildly affected relative to wild type under Pi sufficient condition, similar to previous observation(Arribas-Hernández et al., 2020), root growth was more strongly inhibited upon Pi deficiency (Fig. 3b and Fig. 3c). Expression of *ECT4* under its native promoter sufficed to rescue the triple mutant phenotype, supporting a redundant role for *ECT2, ECT3* and *ECT4* in root growth in response to Pi availability. Null mutants for m^6^A writers are embryonically lethal, hence we used a hypomorphic allele of *FIP7* (*fip37-4*) with a reported ∼10-fold reduction in m^6^A accumulation(Ružicka et al., 2017). Lower levels of m^6^A in *fip37-4* inhibited primary root growth more under Pi-sufficient than Pi-deficient conditions (Fig. 3d). Taken together, these results evidence a role for the m^6^A pathway in Pi homeostasis and primary root growth. Interestingly, ectopic expression of both *ECT4* full-length fused to HA tag (native 3’UTR absent) and *ALT*_*ECT4*_ produced plants with longer primary root in Pi-sufficient condition, while wild type-like growth was observed under Pi-deficient conditions (Fig. 3e). Importantly, *ECT4* expression was unchanged in lines overexpressing *ALT*_*ECT4*_ (Fig. 3f), and overexpression of *ECT4* resulted in similar protein levels in roots grown under low or high Pi level (Fig. 3g). We thus reasoned that *ALT*_*ECT4*_ and *ECT4* might be involved in similar molecular process(es) in primary root development and response to Pi deficiency.

**Figure 3.**
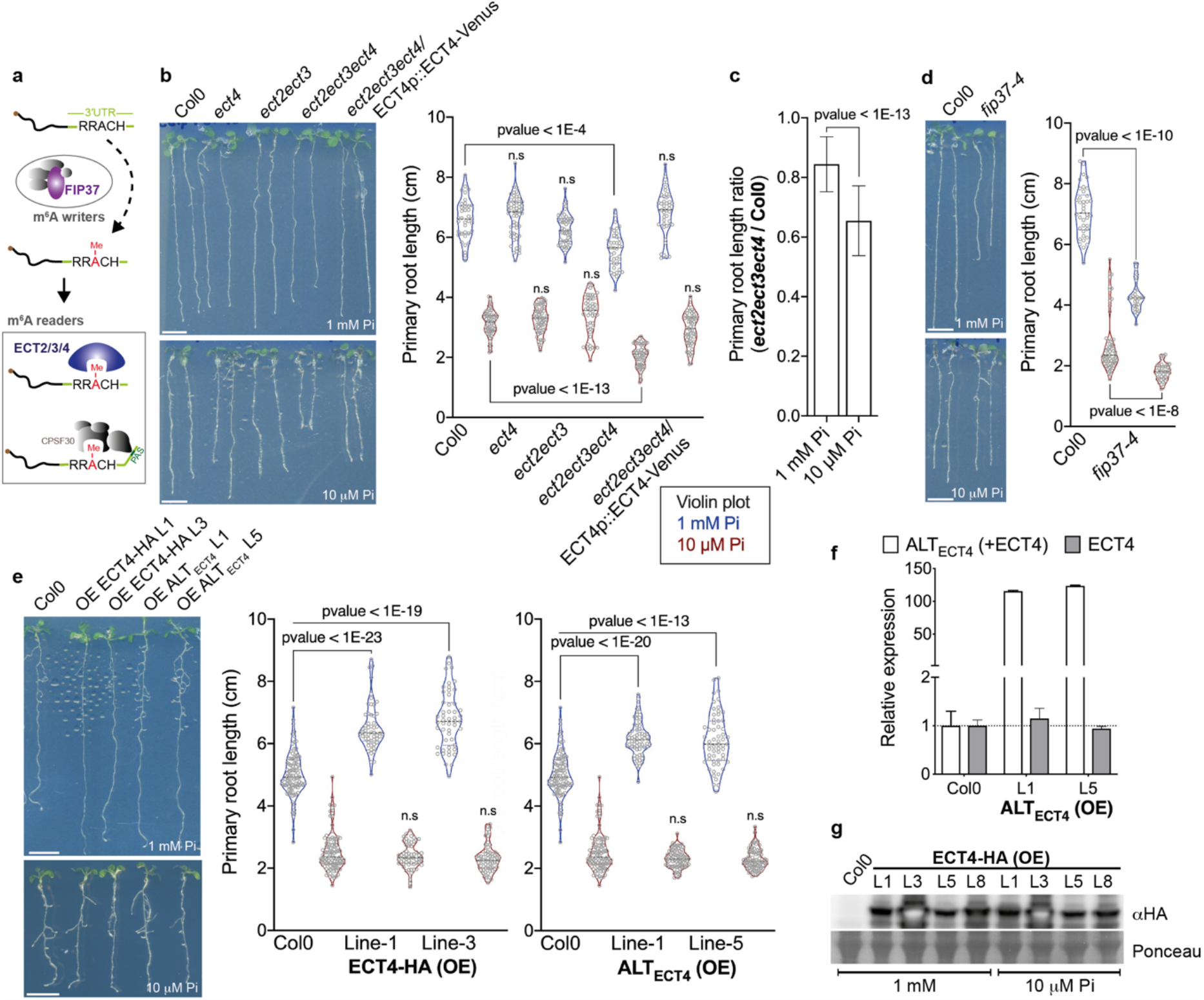
ECT m^6^A readers, FIP37 and *ALT*_*ECT4*_ are involved in primary root development in response to Pi deficiency. **a**, Simplified schematic of RNA modification by the m^6^A pathway. Note that m^6^A erasers are not shown. **b, d** and **e**, Primary root length of plants grown on MS plates for 10 days. Representative plants of each genotype (left panel) and measurements (right panel). One-way ANOVA (adjusted p-value shown; n > 30 biological replicates). *ect2ect3ect4*/*ECT4p::ECT4-Venus*, triple mutant complemented with *ECT4-Venus* (native 3’UTR present) under control of *ECT4* native promoter. *ECT4-HA* (OE) and *ALT*_*ECT4*_ (OE), overexpression in Col0 wildtype background. **c**, Primary root length ratio of *ect2ect3ect4* triple mutant and wild-type in control (1 mM Pi) and Pi deficiency (10 μM Pi) (t-test, n > 30 bio. rep.). Violin plot lines (**b, d** and **e**) are coloured according to treatment (blue, 1 mM Pi control; red, 10 μM Pi deficiency). **f**, RT-qPCR analysis of *ECT4* and *ALT*_*ECT4*_ transcript levels in transgenic lines overexpressing *ALT*_*ECT4*_. Expression relative to Col0 wild-type plants. **g**, ECT4-HA protein fusion accumulation in transgenic lines overexpressing *ECT4-HA* (native 3’UTR absent).

### *ALT*_*ECT4*_ counteracts miR826b-guided translation inhibition of *ECT4*

Analysis of transgenic plants expressing an in-frame *ECT4*-Venus gene fusion (native *ECT4* 3’UTR present) under control of its native promoter, and ECT4-GFP fusion without ECT4 promoter (i.e. GFP driven by *ALT*_*ECT4*_ promoter), showed strong accumulation of free Venus/GFP (detected with anti-GFP antibody) upon Pi starvation (Fig. 4a). In plants expressing *ECT4*-Venus from the *ECT4* promoter, we also observed a reduction in ECT4-Venus protein levels upon Pi starvation, even though their mRNA levels were slightly higher (Fig. S3), suggesting that *ECT4* is posttranscriptionally regulated in Pi-starved plants. Pi and nitrogen homeostasis are entangled processes(Medici et al., 2019). We then assessed whether similar regulation occurs under nitrogen availability. ECT4-Venus protein accumulation was strongly reduced upon nitrogen deficiency (Fig. 4b), without corresponding changes in mRNA levels (Fig. 4d). Nitrogen limitation also did not induce the synthesis of free Venus/GFP, indicating that *ALT*_*ECT4*_ promoter activity was not induced under these conditions, a conclusion also supported by qRT-PCR analysis (Fig. 4d and Fig. S4). Hence, *ECT4* appears to be under similar posttranscriptional regulation in response to both nitrogen and Pi availability, prompting us to search for miRNA target sites within its transcript. Out of predicted target sites (Table S3), a site for miR826b showed parallels with our observations. This miRNA is induced by nitrogen deficiency to trigger miRNA-guided cleavage of glucosinolate synthesis-associated mRNAs, such as AOP2(He et al., 2014). miR826b target site is located at *ECT4* 3’UTR, thus also present in *ALT*_*ECT4*_ transcript. Importantly, *ECT4*/*ALT*_*ECT4*_—miR826b binding exhibits a critical mismatch at nucleotide position 11 of miR826b (Fig. 4c), consistent with miRNA-guided translation inhibition, as such mismatch impairs the ability to guide cleavage(Mallory et al., 2004). In Pi-deficient condition, miR826b, *ECT4* and *ALT*_*ECT4*_ were found at elevated levels, compared to control, while *AOP2* levels were reduced (Fig. 4d). In nitrogen deficiency, however, while *ECT4*’s levels increased to levels similar to that upon Pi deficiency, *ALT*_*ECT4*_ levels were unchanged, miR826b accumulated to much higher levels, and *AOP2* was strongly repressed. Interestingly, transgenic lines overexpressing *ALT*_*ECT4*_ showed weak accumulation of miR826b in phosphate and nitrogen starvation, and its cleavage target *AOP2* showed no reduction in these conditions, suggesting that *ALT*_*ECT4*_ inhibits miR826b accumulation and activity. The observation that strong expression of miR826b under nitrogen deficiency was not associated with a decrease in ECT4 or *ALT*_*ECT4*_ levels indicated that these transcripts are insensitive to miR826b levels. Thus, *ECT4*-miR826b putative interaction might not trigger miR826b-guided cleavage, as opposed to what has been observed with *AOP2* mRNA here and elsewhere(He et al., 2014). We then hypothesized that miR826b guides translation inhibition of *ECT4* mRNA.

**Figure 4.**
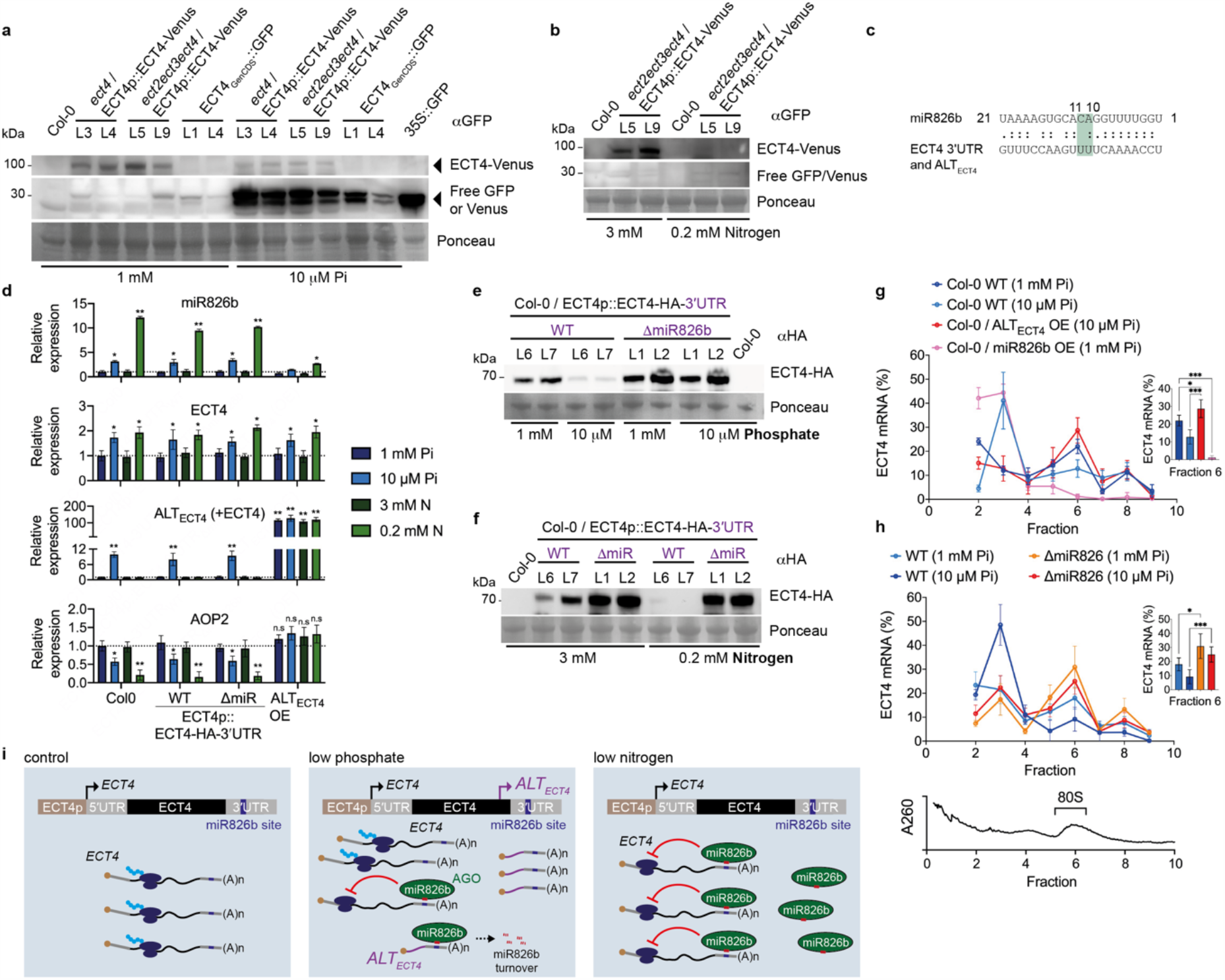
*ALT*_*ECT4*_ counteracts miR826b-guided translation inhibition of *ECT4*. **a** and **b**, *ECT4-Venus* and *ALT*_*ECT4*_ promoter-driven (ECT4_GenCDS_) GFP protein accumulation in Pi (**a**) and nitrogen (**b**) deprivation, in 10-day and 8-day old seedlings, respectively. **c**, Base-pairing of miR826b with *ECT4* 3’UTR and corresponding sequence in *ALT*_*ECT4*_. miR826b nucleotide positions 10 and 11 is highlighted. **d**, RT-qPCR analysis of miR826b (stem loop), *ECT4, ALT*_*ECT4*_ and *AOP2* expression in wild-type and transgenic seedling grown for 10 days (Pi) or 8 days (N). Expression relative to Col0 wild-type plants (t-test, p-value < 0.05* or < 0.01**, n = 3 bio. rep.). **e** and **f**, Representative western blot results for transgenic lines expressing *ECT4-HA* fusion harbouring (WT) or not (ΔmiR826b) the miR826b site (n = 3 bio. rep.). Same seedling pools were used as samples for analysis in **d, e** and **f. g-h**, Distribution of *ECT4* transcript across fractions from sucrose density gradient. Percentage of *ECT4*, relative to total *ECT4* transcript, in fraction 6 (i.e. 80S-bound) is shown (inset bar plot). **h**, Transgenic *ECT4* transcript distribution in Col-0/*ECT4p::ECT4*-HA-3’UTR WT and ΔmiR826b. *ECT4* expression normalized to *ACT2* (t-test, p-value < 0.05* or < 0.001***, n = 3 bio. rep.). Total RNA profile (absorbance at 260 nm) across fractions (lower panel) showing the location of 80S ribosomal fraction. **i**, Proposed model for *ECT4*-miR826b-*ALT*_*ECT4*_ regulatory network. *ECT4* and *ALT*_*ECT4*_ depicted in the *ECT4* genomic locus is shown in the upper part of each panel. Relative to control conditions (left), growth of plants under Pi-deficient condition (middle) induces high expression of the *ALT*_*ECT4*_ transcript, which hijacks miR826b-loaded argonaute(s) (AGO) away from *ECT4* transcript, partially counteracting miR826b-guided translation inhibition of *ECT4* and leading to miR826b turnover. Under low nitrogen, the high level of miR826b is not counteracted by *ALT*_*ECT4*_ expression, resulting in strong inhibition of *ECT4* translation.

Constructs containing the *ECT4* locus, from promoter region to 3’UTR, modified to include a HA tag fused in frame at the C-terminus of ECT4, and either containing or lacking the miR826b site (ΔmiR826b), were expressed in wild type plants. Expression of *ECT4, ALT*_*ECT4*_, miR826b and *AOP2*, were similar to those in non-transgenic plants (Fig. 4d). Under Pi deficiency, lines expressing wild type sequence with the miR826b site showed lower ECT4 protein accumulation (Fig. 4e), albeit its mRNA levels were slightly increased (Fig. 4d). In agreement with our hypothesis of miR826b-guided translation inhibition, ECT4 protein accumulation was higher in ΔmiR826b lines, under either Pi-sufficient or deficient conditions. Under nitrogen deficient condition, in which miR826b induction was the highest, ECT4 protein was barely detected in wild type lines, while its accumulation was high in the ΔmiR826b lines (Fig. 4f). Similarly, under nitrogen sufficient conditions, the level of ECT4 protein was higher in the ΔmiR826b lines compared to the wild type background (Fig. 4f). To assess whether protein accumulation was a direct effect of translational control, we analysed the distribution of *ECT4* transcript across fractions in a sucrose gradient (Fig. 4g-h). Endogenous *ECT4* transcript was found less associated with 80S ribosome in non-transgenic plants under low Pi (Fig. 4g). Overexpression of *ALT*_*ECT4*_ resulted in increased *ECT4* association with 80S under low Pi. Strikingly, overexpression of miR826b virtually abolished *ECT4* association with 80S in plant grown under normal Pi conditions, in which in non-transgenic plants *ECT4*-80S was elevated. We then analysed *ECT4* in transgenic lines expressing the WT or mutant (ΔmiR826b) version for the miRNA target site (Fig. 4h). The effect of Pi starvation in these lines evidenced a clear negative correlation between low Pi and the presence of the miR826b target site. In ΔmiR826b plants, Pi starvation had no effect on ECT4 association with 80S, whereas in the presence of miR826 binding site resulted in low 80S association in plants grown under low Pi. These results demonstrate that *ECT4* is targeted by miR826b in a translation inhibition manner, and this regulation is less pronounced in Pi-starved than in N-starved plants, possibly because *ALT*_*ECT4*_ levels are induced by Pi deficiency but not by nitrogen.

*ALT*_*ECT4*_ antagonism of miR826b accumulation and, hence, the reduced miRNA inhibition of its targets (i.e. *AOP2* transcript cleavage and *ECT4* translation inhibition), is consistent with *ALT*_*ECT4*_ been recognized by miR826b as target mimic. To date, target mimicry has only been found in *trans*, i.e. a noncoding RNA harbouring miRNA target site(s) with mismatch(es) at the expected cleavage site, between positions 9 and 11, is expressed from a locus other than the miRNA targets, such as in the IPS1-miR399-PHO2 network(Franco-Zorrilla et al., 2007). In our proposed model, *ECT4* is under translation inhibition guided by miR826b in the absence or low levels of *ALT*_*ECT4*_ (Fig. 4i). In Pi starvation, however, *ALT*_*ECT4*_ induction provides an uncleavable noncoding target for miR826b-loaded argonaute (AGO), sequestering miR826b and leading to its turnover, similar to IPS1 sequestration of miR399.

Our work revealed several alternative TSS tightly controlled by Pi availability and, in particular, this process regulates the m^6^A reader *ECT4*. Although the use of alternative TSS is a widespread process in plants, current studies have been focused almost exclusively on transcripts isoforms that retain the coding sequence of the reference transcript(Kurihara et al., 2018; Ushijima et al., 2017; Wang et al., 2019). The discovery of ECT4-miR826b-ALT_ECT4_ regulatory network evidence a broader role for alternative TSS, beyond the synthesis of protein isoforms, in controlling the expression of the main protein-coding transcript. We envisage that such mode of action might have been an important evolutionary pressure for alternative TSS promoter formation.

## Methods

### Plant material and constructs

All Arabidopsis plants used in this study, including mutants and transgenic plants, were in the Columbia (Col0) background. T-DNA insertion lines *ect4-2* (*ect4*), *ect2-1*/*ect3-1* (*ect2ect3*) and *ect2-1*/*ect3-1*/*ect4-2* (*ect2ect3ect4*), as well as transgenic lines *ect4*/ECT4p::ECT4-Venus and *ect2ect3ect4*/ECT4p::ECT4-Venus were gifts from Peter Brodersen and were previously described(Arribas-Hernández et al., 2018). *fip37-4* (SALK_018636) was obtained from Nottingham Arabidopsis Stock Centre (NASC). *ECT4* promoter region (880 upstream of *ECT4* TSS(Arribas-Hernández et al., 2018)), *ECT4* and *ALT*_*ECT4*_ sequences were amplified from genomic DNA and cloned into an ENTR vector (Table S4). Binary constructs ECT4pro::GUS and ALT_ECT4_pro::GUS were produced by LR cloning using pMDC163(Curtis and Grossniklaus, 2003), and ALT_ECT4_pro::GFP by LR using pMDC107(Curtis and Grossniklaus, 2003). ECT4p::ECT4-HA-3’UTR was produced using the forward oligo for ECT4 promoter and reverse for ECT4 3’UTR, cloned into an ENTR vector, followed by overlapping PCR to add an in-frame HA tag. Overlapping PCR was also used to remove the miR826b site in ΔmiR826b construct. The final ENTR vectors were recombined with LR cloning into pFAST-R01(Shimada et al., 2010). RTNLB1 and ALT_RTNLB1_ 5’UTR were amplified from genomic DNA (Table S4), cloned into an ENTR vector and LR recombined with a protoplast-expression construct previously described(Deforges et al., 2019b).

### Plant growth conditions and treatments

Plants were grown on half-strength Murashige and Skoog (MS) salts (Duchefa M0255) containing 1% (w/v) sucrose and 0.7% (w/ v) agar, and cultivated at 22^0^C under continuous light. For experiments with variable Pi concentration, MS salts without Pi (Caisson Labs, MSP11) were supplemented with Pi buffer (pH5.7, 93.5% w/v KH_2_PO_4_ and 6.5% w/v K_2_HPO_4_) to reach a final concentration of 1 mM or 10 uM. For nitrogen starvation, MS salts without nitrogen (Caisson Labs, MSP07) were supplemented with nitrogen to a final concentration of 3 mM (1 mM NH_4_NO_3_ and 1 mM KNO_3_) or 0.2 mM (0.07 mM NH_4_NO_3_, 0.07 mM KNO_3_ and 0.93 mM KCl). Sample material produced for RNA-Seq and PacBio have been previously described(Deforges et al., 2019a). In brief, wild type Col0 seeds were germinated in half-strength MS liquid medium containing 1 mM or 0.1 mM Pi using MS salts without Pi, as described above. Pi-deficient plants were harvested for total RNA extraction 7 days after germination. Hormone treatments were performed in standard half-strength MS medium, as above. Ten days after germination, wild type Col0 seedlings were flooded with a solution of half-strength MS containing 5 uM indole acetic acid (IAA), 10 uM abscisic acid (ABA), 10 uM methyl jasmonate (MeJA), or no hormone for the untreated control. After 3 h of incubation, roots and shoots were harvested separately (3 independent biological replicates).

### RNA-Seq and JunctionSeq analysis

The RNA-Seq dataset has been previously published(Deforges et al., 2019a). Here, we applied JunctionSeq analysis(Hartley and Mullikin, 2016) to all RNA-Seq reads mapped against TAIR10 annotated genes. JunctionSeq was set to analyze differential exons only, thus splice junctions were excluded, with false discovery rate (FDR) of 0.01.

### TeloPrime cDNA library preparation with cap-dependent linker ligation

The same RNA extracted from plants were used for RNA-Seq (see above) and PacBio long read analysis. The TeloPrime Full-Length cDNA Amplification Kit (Lexogen, Austria) was used for generating full-length cDNA from 1 μg of total RNA, by the Genomic Technology Facility at the University of Lausanne. Briefly, first strand cDNA synthesis is initiated by a 3’ oligo-dT anchoring primer (RP: 5’-TCTCAGGCGTTTTTTTTTTTTTTTTTT-3’) and reverse transcription. The cDNA:RNA hybrid molecules were column purified and ligated to the double-stranded linker (cap-dependent linker ligation, CDLL) carrying a 5’ C overhang, thus allowing base pairing with the G nucleotide of the 5’ mRNA CAP. Ligation products were again column purified and the resulting eluted fragments were converted to full-length double-stranded cDNA by second strand cDNA synthesis.

### PacBio library preparation and sequencing

PacBio library preparation and sequencing in a Pacific Biosciences Sequel system, were performed by the Genomic Technology Facility at the University of Lausanne.

### PacBio long reads and alternative TSS transcript analysis

PacBio consensus sequences were obtained from subreads using css v4.1.0 (https://github.com/PacificBiosciences/ccs). Since the first strand cDNA synthesis is initiated by a 3’ oligo-dT anchoring primer (RP:5’-TCTCAGGCGTTTTTTTTTTTTTTTTTT-3’), the orientation of the consensus sequence was established by detecting the presence of an “AGGCGTTTT” sequence within the first 50 nucleotides of the consensus (indicating it should be reversed complemented) or the presence of a “AAAACGCCT” within the last 50 nucleotides of the consensus sequence (indicating it was properly oriented). Full length sequences were selected by detecting the presence of a “CTCACTATAG” within the first 100 nt of oriented sequences. Full-length sequences were mapped against the Arabidopsis genome TAIR10 using minimap2 v2.17-r954-dirty using the option -ax splice:hq --seed 11 -t24 --secondary=no -r300k. Forward and reverse oriented sequences were used to detect potential alternative transcription start sites between Pi starvation and control conditions. Read number for full-length transcripts (annotated full-length, truncated and exon skipped exon) were compared using chi-squared test (uncorrected for low counts or FDR) were reported in Table S1.

### RNA isolation and RT-qPCR

Total RNA for RT-qPCR analysis was extracted from root, shoot and seedling using an RNA purification kit, as described by the manufacture (Jena Bioscience, PP-210), followed by DNase I treatment. cDNA was synthesized from 0.5 ug RNA using M-MLV Reverse Transcriptase (Promega, M3681) and oligo d(T)15 following the manufacturer’s instructions. qPCR analysis was performed using SYBR select Master Mix (Applied Biosystems, 4472908) with primer pairs specific to genes of interest and ACT2, used for data normalization. Primer sequences are listed in Table S4. RT-qPCR for miR826b was performed as previously described(Varkonyi-Gasic et al., 2007). Briefly, 1 ug of total RNA extracted with Trizol was used for stem loop reverse RT-qPCR using primers listed in Table S4. snoR101 was used to normalize the miR826b accumulation.

### Polysome fractionation

Arabidopsis seedlings were frozen, ground and the polysomes were extracted essentially as previously described (Mustroph et al., 2009) with minor modifications. Briefly, the powder was resuspended in 2 volumes of polysome extraction buffer [200 mM Tris pH 9.0, 200 mM KCl, 1 % (w/v) deoxycholate, 25mM EGTA, 1 % (v/v) detergent mix containing equal proportion of Brij-35, Triton X-100, octylphenyl-polyethylene glycol and Tween-20, 1 % (w/v) polyoxyethylene 10 tridecyl ether, 35 mM MgCl2, 1 mM DTT and 100 ug/ml cyclohexomide], and incubated on ice for 15 min. The mixture was then centrifuged at 4°C for 15 min at 20,000 g, and the supernatant was layered on top of a 10 mL sucrose cushion [60 % (w/v) sucrose, 400 mM Tris pH 9.0, 200 mM KCl, 25 mM EGTA, 35 mM MgCl2, 1 mM DTT and 100 ug/mL cyclohexomide] and centrifuged for 3 h at 170,000 xg. The pellet was resuspended in 100 uL of the same sucrose cushion buffer without sucrose, and loaded on top of 5 mL 15-60 % continuous sucrose gradients. The gradients were centrifuged for 75 min at 237,000 g in swinging buckets and fractionated using a gradient holder apparatus (Brandel) into 12 fractions. During gradient fractionation, the UV absorbance was continuously monitored in order to detect the position of the different complexes within the gradient. RNA was extracted from each collected fraction using the RNA Clean and Concentrator-25 kit following the manufacture’s instruction (Zymo Research, R1017). Reverse transcription and qPCR were performed as described above. Polysome association was calculated as relative proportion of mRNA in each fraction of the gradient, as previously described (Faye et al., 2014).

### uORF analysis and domain and signal peptide prediction

For each selected transcript, one transcript sequence (Araport11) derived from a corresponding representative gene model was obtained via Sequence Bulk Download from TAIR. Upstream open reading frames (uORFs) were defined as any ORF of less than or equal to 200 nt in length, in which its ATG START codon was located at a position less than or equal to 200 nt upstream the main ORF ATG. We then translated the longest, reference ORF within each transcript and used InterProScan 5(Jones et al., 2014) to detect and predict functional domains (default parameters). Similarly, each reference amino acid sequence was used to predict signal peptide using TargetP-2.0(Almagro Armenteros et al., 2019) (organism group, plant). For alternative transcripts, the longest ORF ending at the corresponding reference ORF’s STOP codon, was selected for comparative analysis.

### miRNA target site prediction

miRNA target sites within ECT4 transcript sequence were predicted using psRNATarget(Dai and Zhao, 2011). *A. thaliana* preloaded small RNAs (miRbase release 21) were applied with a maximum expectation set to 5 (low stringency) and range of central mismatch leading to translational inhibition set to 9-11 nt (default). ECT4 genomic sequence with predicted miRNA target sites are found in Table S3.

### Immunoblot analysis

Approximately 50-200 mg of seedling material was ground in liquid nitrogen and dissolved in SDS sample buffer (50 mM Tris-HCl pH 6.8, 2% w/v SDS, 10% v/v glycerol, 1% v/v beta-mercaptoethanol, 12.5 mM EDTA, and 0.02 % w/v bromophenol blue). Total protein was separated in a 4-20% gradient SDS-PAGE gel, transferred to a PVDF membrane, and incubated with corresponding primary and secondary antibodies before chemiluminescence-image capture. For HA-tagged proteins, we used rat anti-HA (Roche, 3F10 clone) at 1:7,000 dilution and goat anti-rat secondary (Abcam, ab7097) at 1:1,500 dilution. For GFP or Venus-tagged proteins, we used rabbit anti-GFP (Abcam, ab6556) at 1:2,500 dilution and goat anti-rabbit (Agrisera, AS09 602) at 1:30,000 dilution.

### Reporter gene expression analysis

To localize GUS activity in transgenic lines expressing promoter-GUS fusion, seedlings were treated as previously described(Jefferson et al., 1987).

## Data availability

PacBio long read data available under the BioProject ID PRJNA649694.

## Acknowledgments

The authors thank Peter Brodersen, University of Copenhagen, for kindly providing us with several transgenic lines. We thank Nicolas Guex and Christian Iseli of the University of Lausanne Bioinformatic Center for PacBio data analysis. PacBio sequencing was performed by the University of Lausanne Genomic Technologies Facility. We also thank Dominique Jacques-Vuarambon for technical assistance. This work was supported by a <u>Swiss National Science Foundation (Schweizerische Nationalfonds)</u> Sinergia grant (CRSII3_154471 to Y.P.).

## Author contributions

RSR, JD and YP conceived and designed the study. JD performed RNA-Seq and JunctionSeq analysis. JC performed GUS staining. RSR performed all other experiments, except for PacBio data analysis (outsourced). RSR and YP wrote the paper. All authors read and approved the final manuscript.

## Competing interests

The authors declare no competing interests.

## Additional information

### Extended data

**Figure S1.**
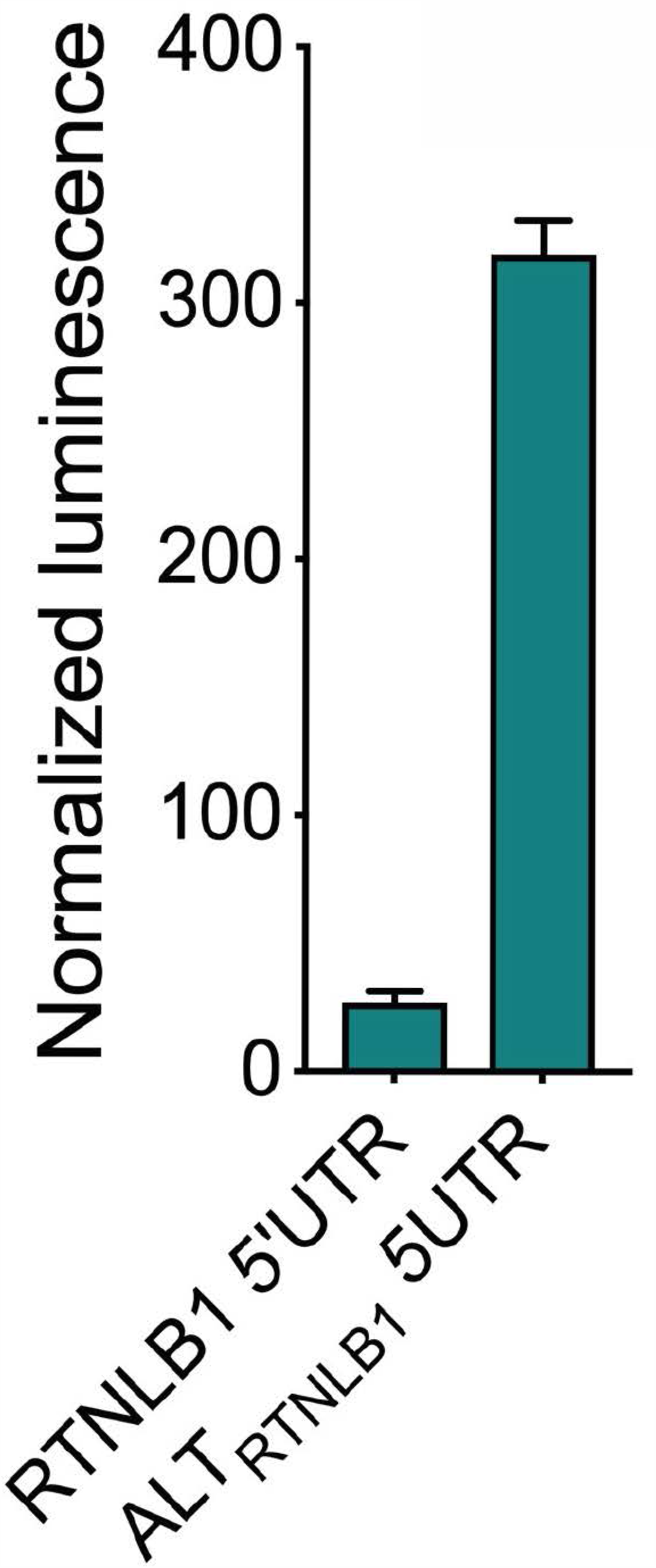
Effect of *RTNLB1* 5’UTR and associated alternative TSS (*ALT*_*RTNLB1*_) 5’UTR on protein accumulation. Normalized luminescence in transfected protoplasts (n = 6 biological replicates). Luminescence values produced in protoplasts from the expression of nano Luciferase (nLuc) fused to either *RTNLB1* 5’UTR or the *ALT*_*RTNLB1*_ 5’UTR were normalized against luminescence from the firefly luciferase (Fluc). Both nLuc and Fluc luciferase genes were expressed from the same plasmid using the PcUBI4-2 and AtRPS5A promoter, respectively.

**Figure S2.**
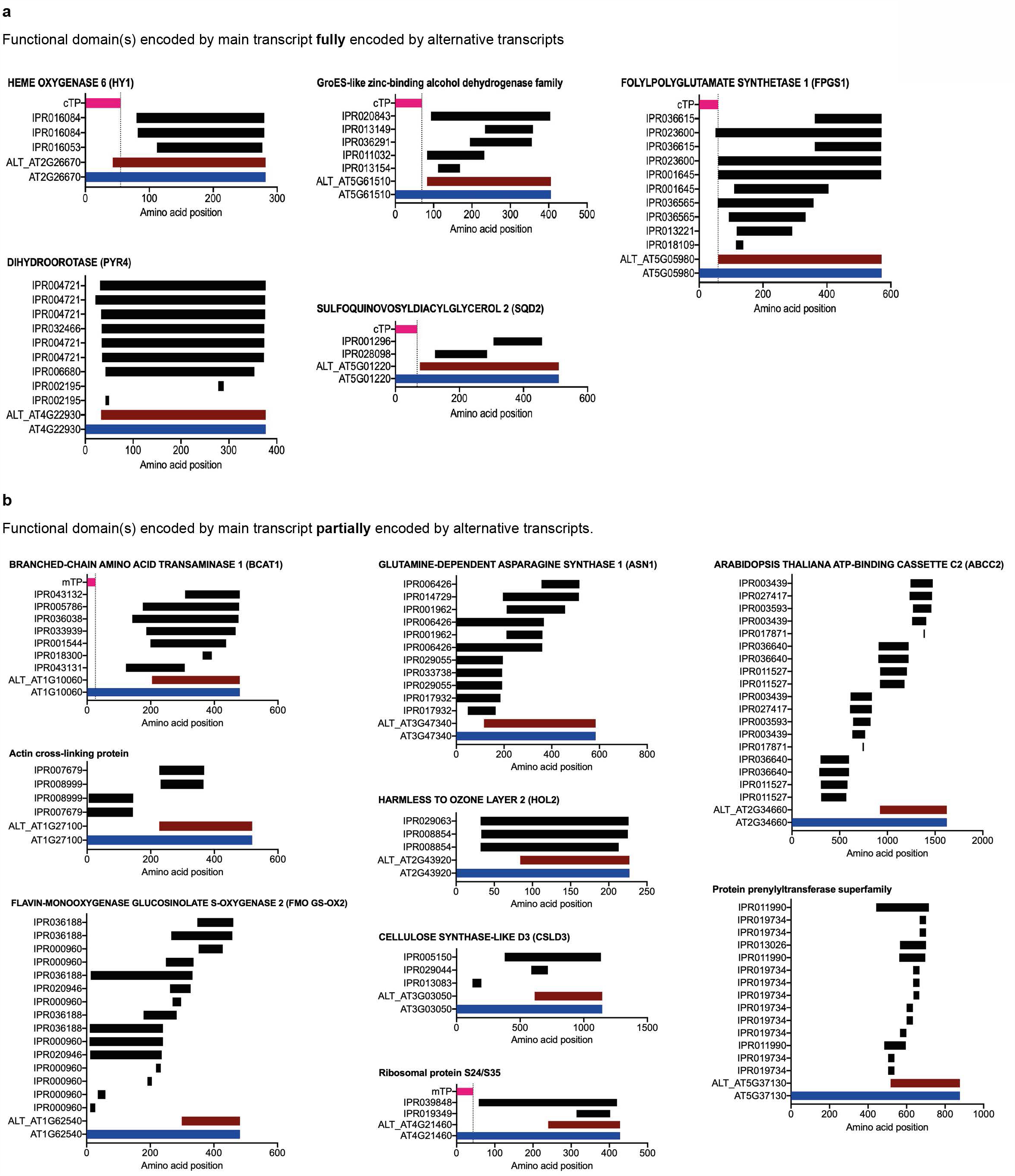
Functional features associated with reference transcript-encoded amino acid sequence. **a**, Proteins with full-length functional domain(s) encoded by reference and alternative transcripts. **b**, Proteins with functional domain(s) encoded by reference and partially by alternative transcripts. Predicted domains using InterProScan depicted as IPR-number (black bar). SP, signal peptide (pink bar). Reference protein (locus name; blue bar) and alternative transcript-encoded amino acid sequence (ALT_locus name; red bar). Alternative transcript-encoded amino acid sequence was defined as the longest ORF ending at the reference transcript’s STOP codon, i.e. exhibiting exactly the same (complete or partial) amino acid sequence as that found in the reference protein.

**Figure S3.**
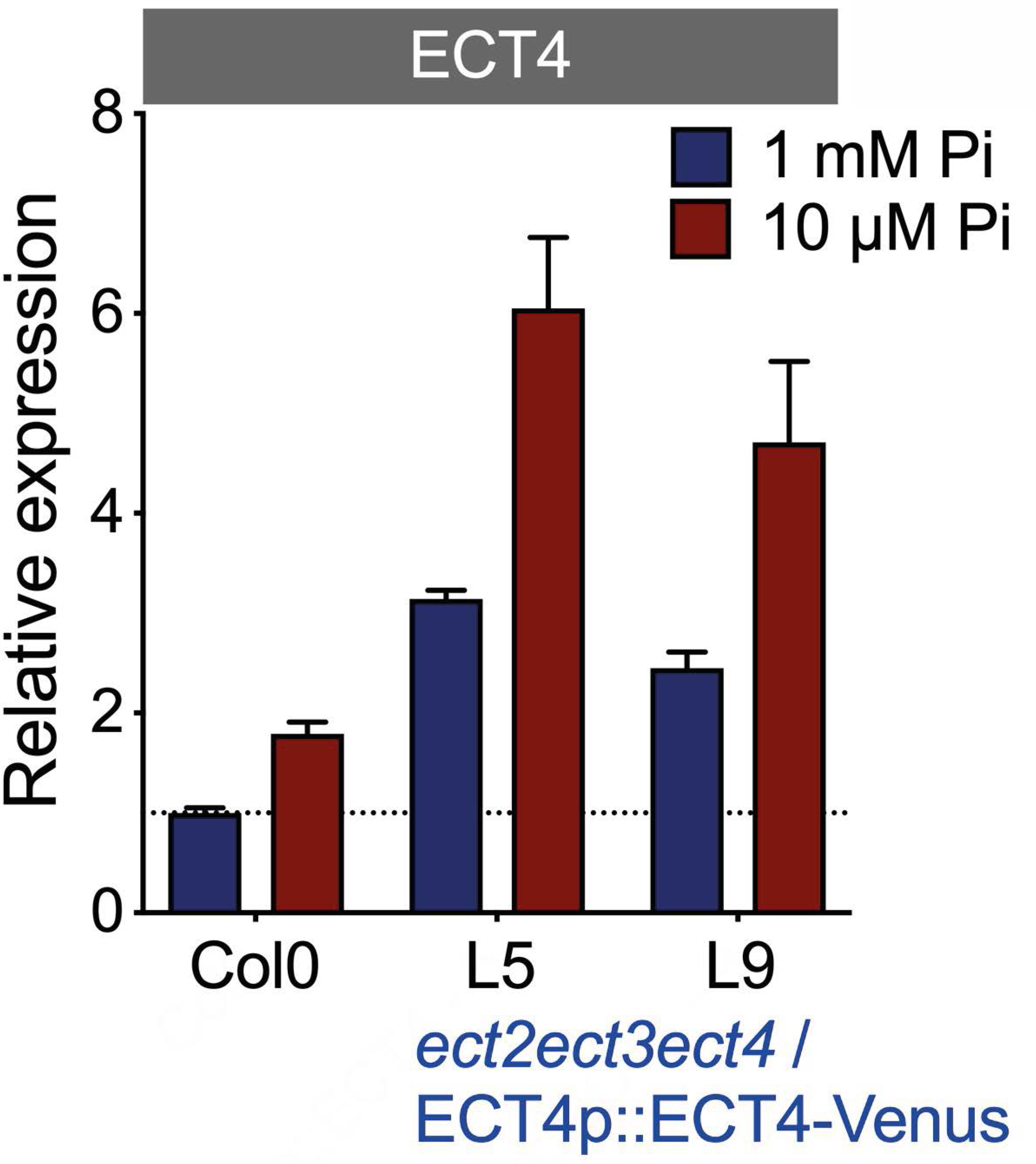
*ECT4* expression in phosphate deprivation. *ECT4* expression profile of Pi-deprived plants grown on MS plates. *ect2ect3ect4*/*ECT4p::ECT4-Venus*, triple mutant complemented with *ECT4-Venus* (native 3’UTR present) under control of *ECT4* native promoter. RT-qPCR using oligo pair specific to *ECT4*, endogenous and transgene (n = 3 bio. rep.).

**Figure S4.**
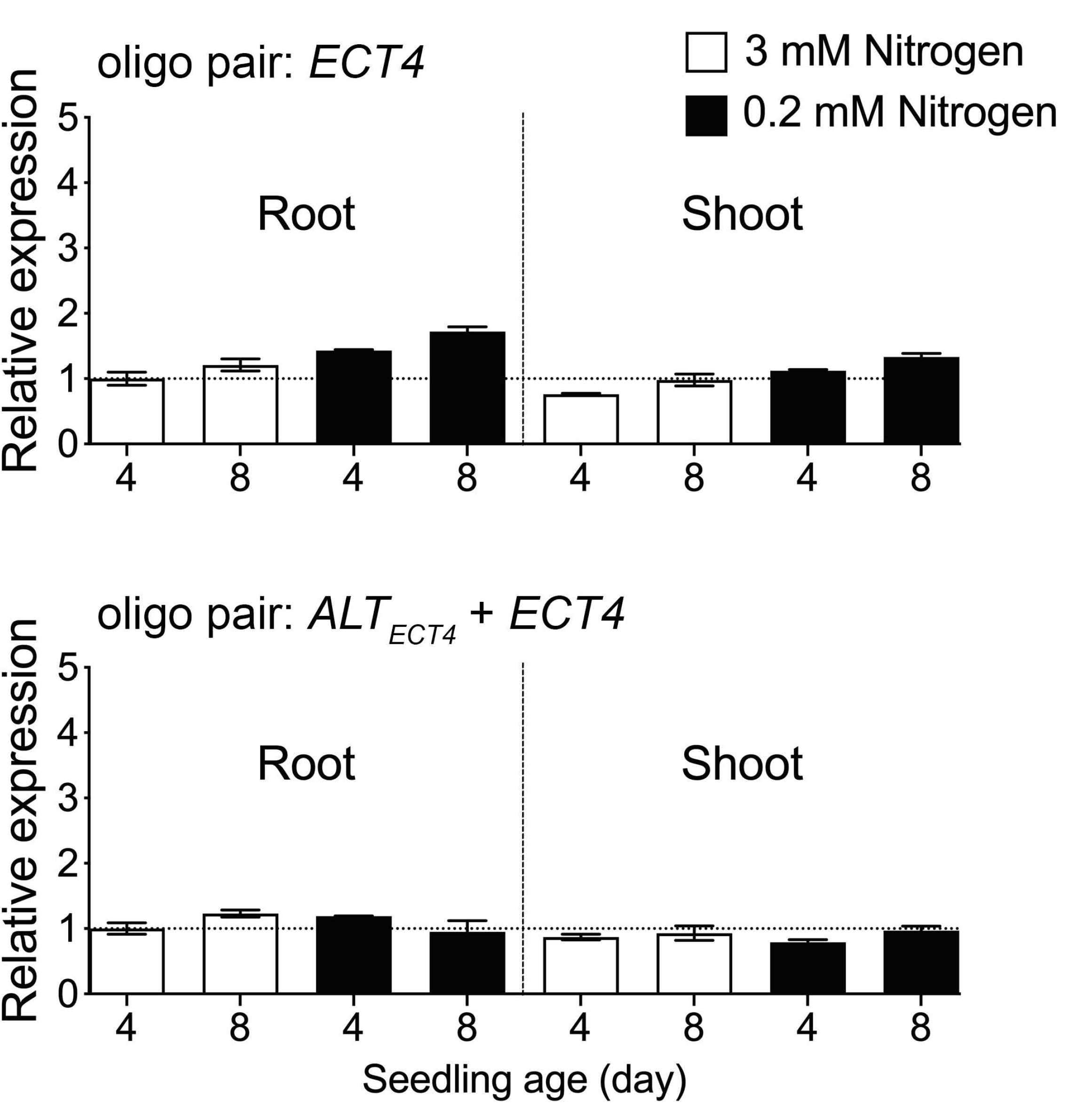
*ECT4* and *ALT*_*ECT4*_ expression in nitrogen deficiency. *ECT4* and *ALT*_*ECT4*_ expression profile in plants grown on nitrogen-deficient media for 4 and 8 days. RT-qPCR using oligo pair (see Figure 2b) specific to *ECT4* (top panel) or specific to both *ECT4* and *ALT*_*ECT4*_ (bottom panel) (n = 3 bio. rep.).

**Table S1**. Alternative TSS genes. Genes identified by JunctionSeq and PacBio, as well as analysis of uORF, signal peptide and domain.

**Table S3**. Predicted miRNA target sites for ECT4. Predicted miRNA target sites annotated on ECT4 genomic sequence, presented in GeneBank format.

**Table S2.**
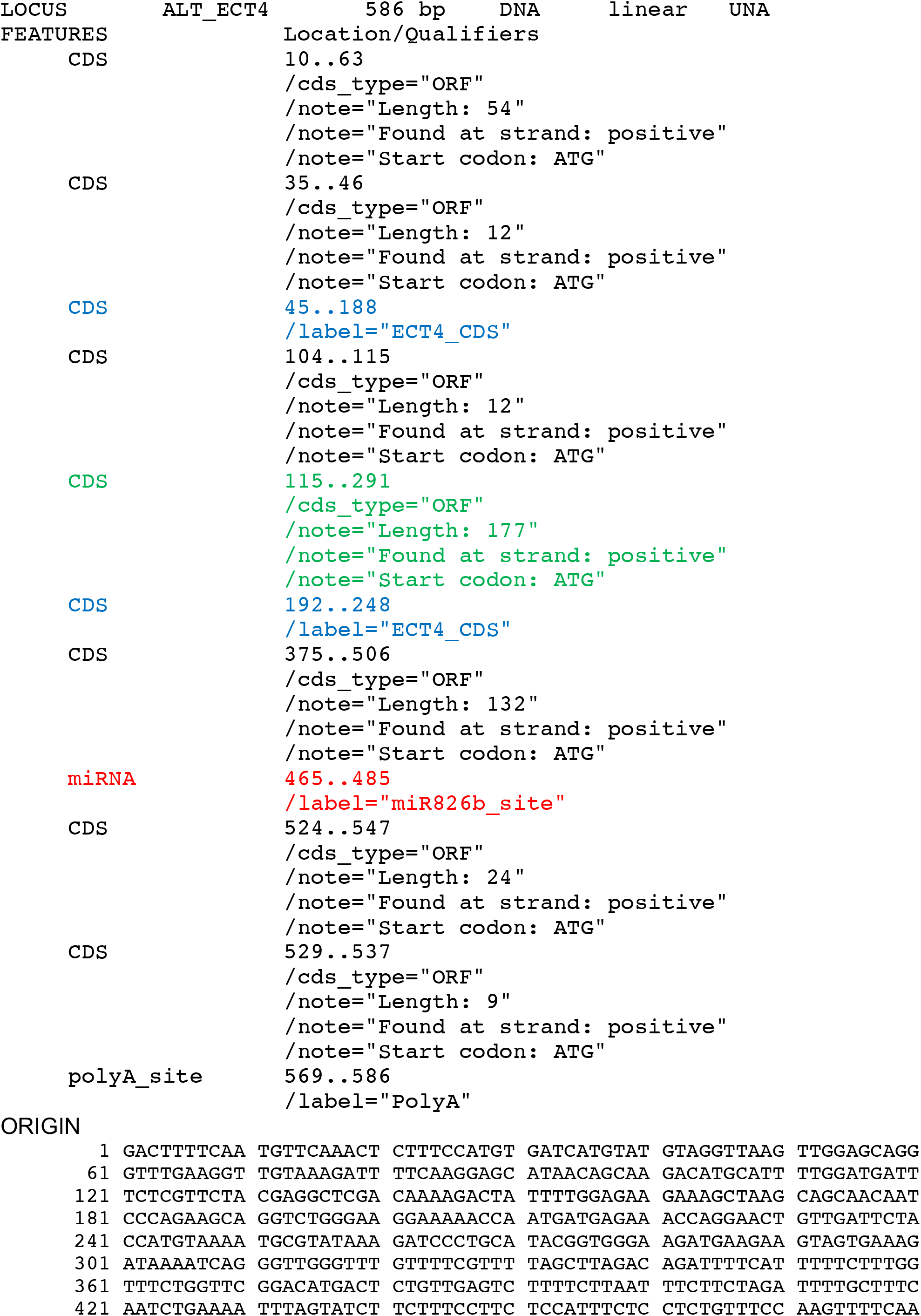
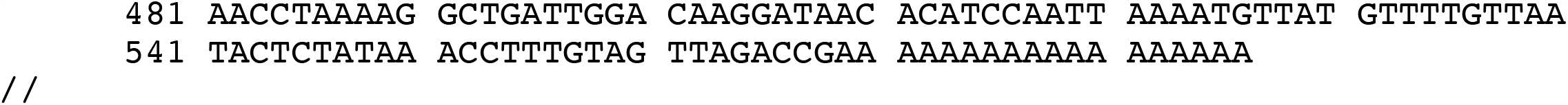
Annotated ALT_ECT4_ sequence. ALT_ECT4_ sequence obtained from PacBio long read and presented in GeneBank format.

**Table S3.**
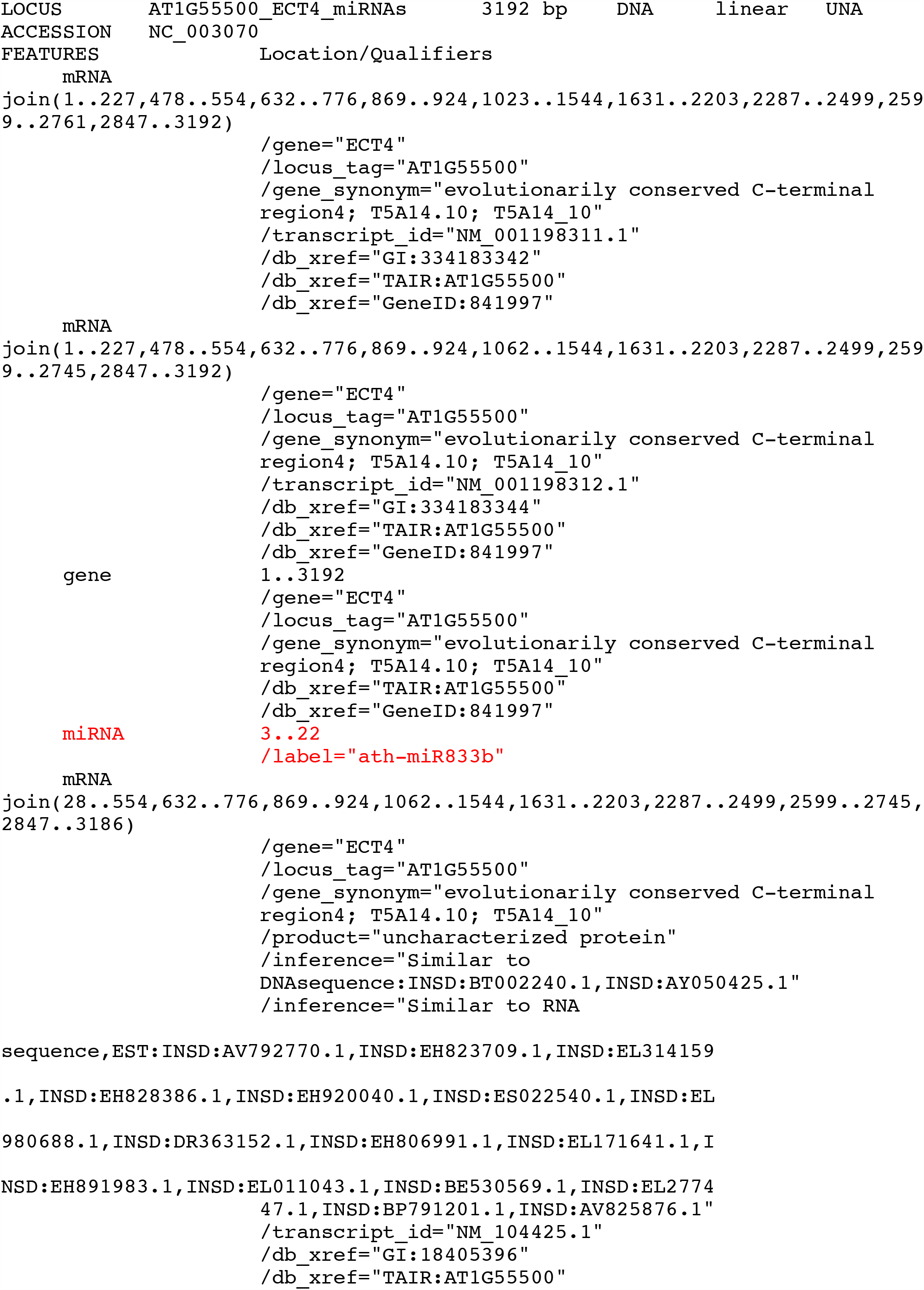
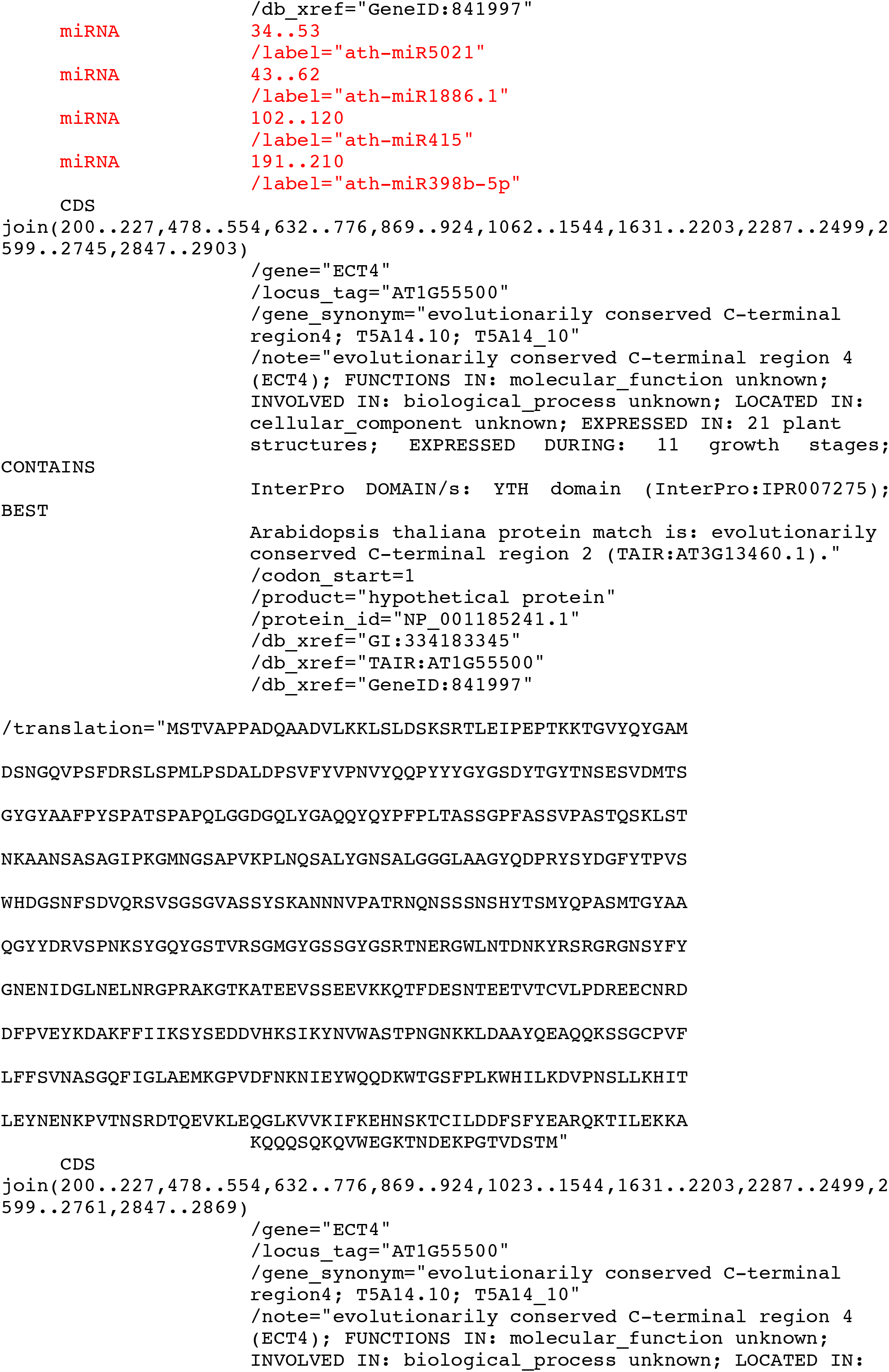
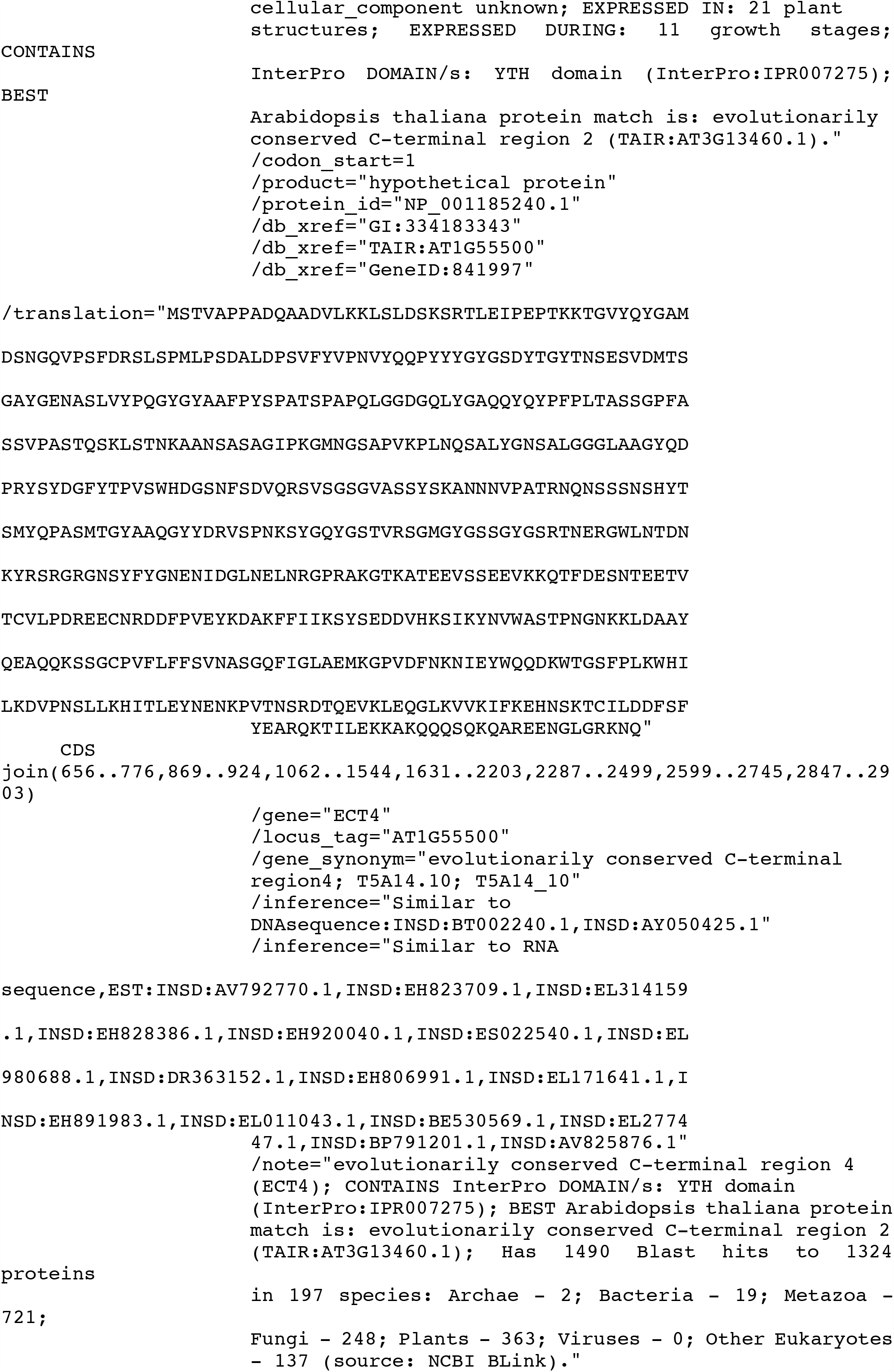
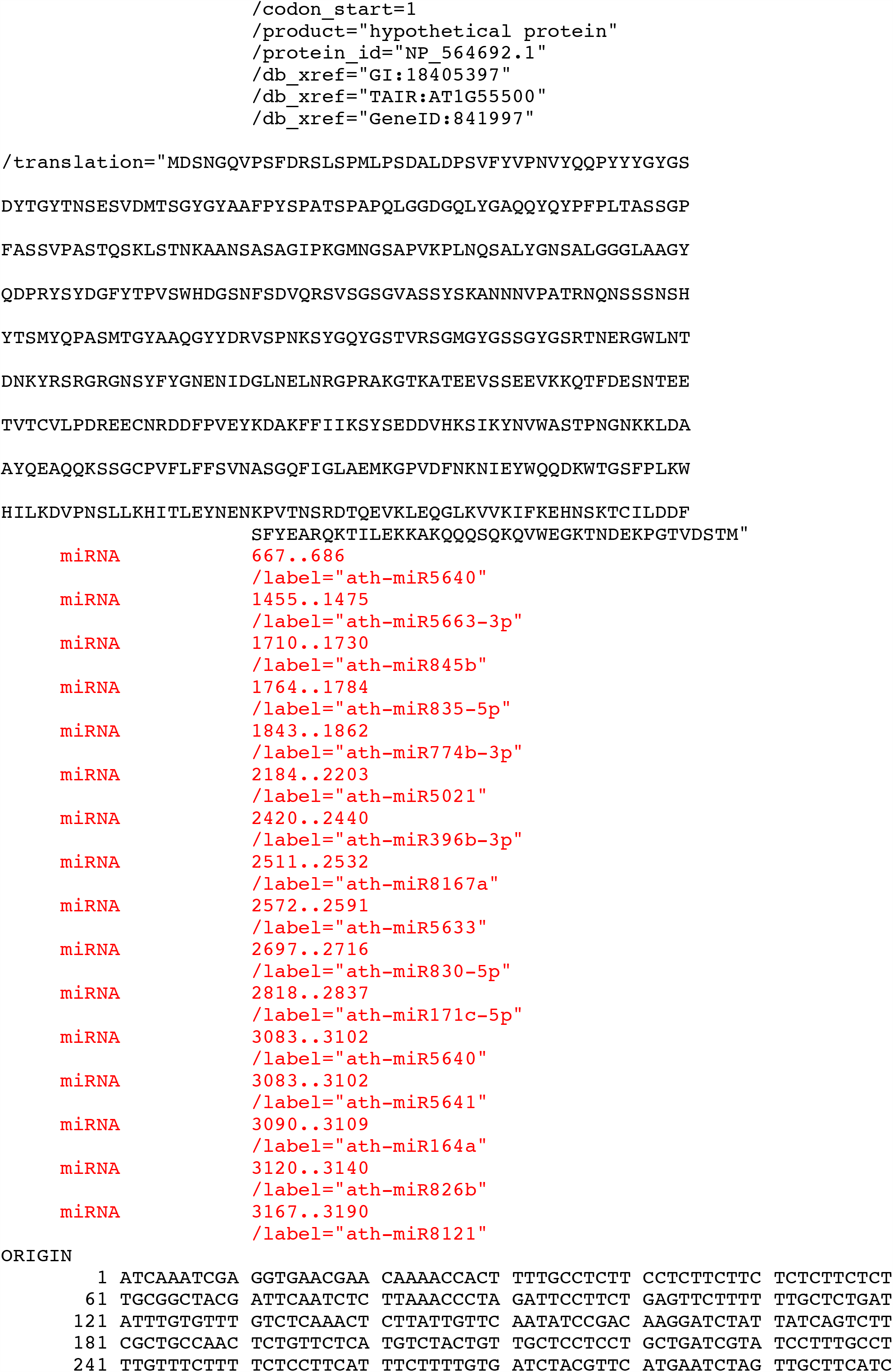
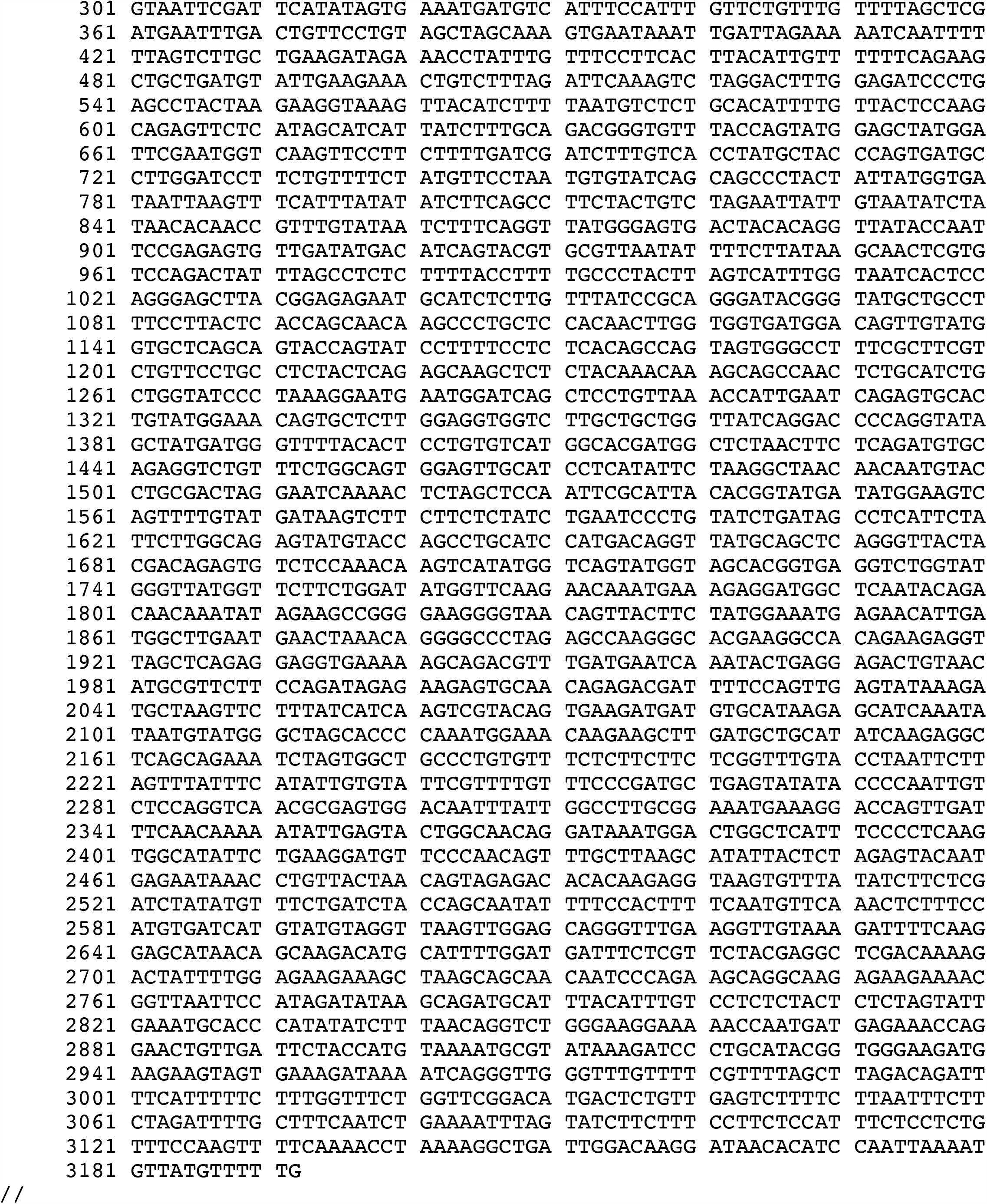
Predicted miRNA target sites for ECT4 (AT1G55500). GeneBank file for predicted miRNA target sites annotated on ECT4 genomic sequence (accession number NC_003070).

**Table S4.**
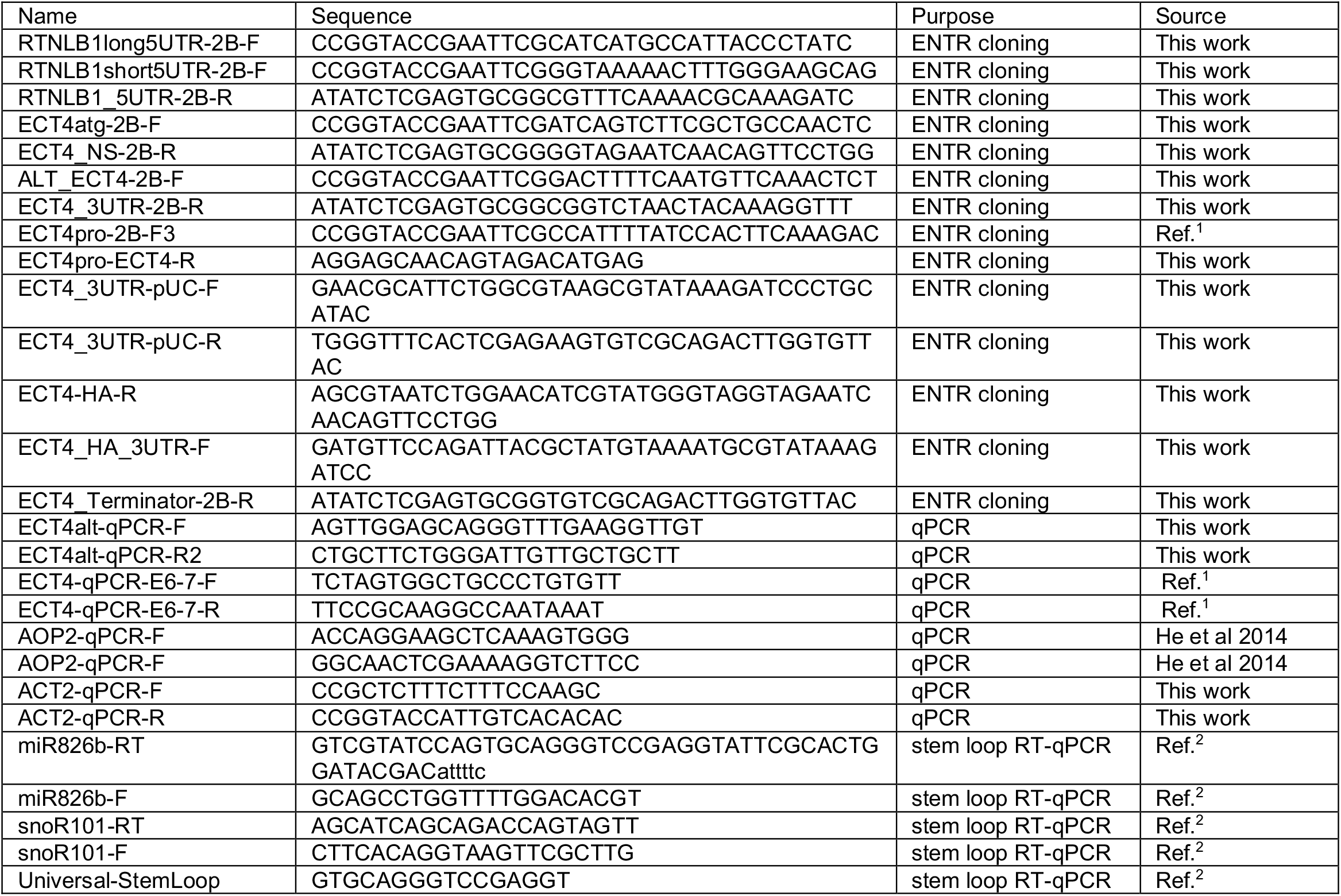
Primer list.

## Notes

### Competing Interest Statement

The authors have declared no competing interest.

### Summary of Updates

#version2 Fixed typos.

